# MixOmics Integration of Biological Datasets Identifies Highly Correlated Key Variables of COVID-19 severity

**DOI:** 10.1101/2023.09.14.557558

**Authors:** Noa C. Harriott, Michael S. Chimenti, Amy L. Ryan

**Affiliations:** Department of Anatomy and Cell Biology, Carver College of Medicine, University of Iowa, Iowa City IA 52240; Department of Stem Cell Biology and Regenerative Medicine, University of Southern California, Los Angeles CA 90033; Hastings Center for Pulmonary Research, Division of Pulmonary, Critical Care and Sleep Medicine, Department of Medicine, University of Southern California, Los Angeles CA 90033; Iowa Institute of Human Genetics, Carver College of Medicine, University of Iowa, Iowa City IA 52240

**Keywords:** Biomarkers, DIABLO, Machine Learning, Multiomics, Proteomics, SARS-CoV-2, Transcriptomics

## Abstract

**Background:** Despite several years since the COVID-19 pandemic was declared, challenges remain in understanding the factors that can predict the severity of COVID-19 disease and complications of SARS-CoV-2 infection. While many large-scale Multiomic datasets have been published, integration of these datasets has the potential to substantially increase the biological insight gained allowing a more complex comprehension of the disease pathogenesis. Such insight may improve our ability to predict disease progression, detect severe cases more rapidly and develop effective therapeutics.

**Methods:** In this study we have applied an innovative machine learning algorithm to delineate COVID-severity based on integration of paired samples of proteomic and transcriptomic data from a small cohort of patients testing positive for SARS-CoV-2 infection with differential disease severity. Targeted plasma proteomics and an onco-immune targeted transcriptomic panel was performed on sequential samples from a cohort of 23 severe, 21 moderate and 10 mild COVID-19 patients. We applied DIABLO, a new integrative method, to identify multi-omics biomarker panels that can discriminate between multiple phenotypic groups, such as the varied severity of disease in COVID-19 patients.

**Results:** As COVID-19 severity is known among our sample group, we can train models using this as the outcome variable and calculate features that are important predictors of severe disease. In this study, we detect highly correlated key variables of severe COVID-19 using transcriptomic discriminant analysis and multi-omics integration methods.

**Conclusions:** This approach highlights the power of data integration from a small cohort of patients offering a better biological understanding of the molecular mechanisms driving COVID-19 severity and an opportunity to improve prediction of disease trajectories and targeted therapeutics.

## Background

Since the COVID-19 pandemic ensued, a plethora of symptoms have been identified to lead to stratification of disease severity within patients infected with SARS-CoV-2. Symptoms are akin to those observed in severe acute respiratory distress syndrome (SARS) inclusive of fever, dry cough, exhaustion, loss of taste and smell and shortness of breath [1–5]. The efficiency of the host’s immune response and the infectivity of the SARS-CoV-2 are two core factors that define disease pathogenesis and viral survival. Despite a vast array of studies investigating the pathogenesis of COVID-19, we still do not fully comprehend the biomarkers that can predict severe disease, nor the biological pathways contributing to disease progression and severity [6–12]. High-throughput ‘Omics technologies have been applied to rapidly understand the mechanistic pathways of viral infection for several viruses, including dengue, zika and West Nile virus [13–15]. Similar large-scale Multiomic studies have been published over the past 3 years investigating the viral pathogenesis of SARS-CoV-2 [16–26].

As stand-alone datasets they provide valuable information on disease pathogenesis. However, integration of these datasets has the potential to substantially increase the depth of biological insight gained. Systems biology approaches can leverage multi-omics datasets, identify molecular biomarkers of disease and capture biological network complexity. Data Integration Analysis for Biomarker Discovery, DIABLO, is an integrative method that can be applied to identify multi-omics biomarker panels that can discriminate between multiple phenotypic groups, such as the varied severity of disease in COVID-19 patients [27, 28]. An in-depth understanding of the biological changes occurring in response to SARS-CoV-2 infection can be assimilated through evaluation of cellular and molecular features including proteins, RNA and DNA. In this study, we detect biomarkers of severe COVID-19 using transcriptomic discriminant analysis and multi-omics integration methods. Since COVID-19 severity is known among our sample group, we can train models using this as the outcome variable and calculate features that are important predictors of severe disease. Our study highlights the power of integrating datasets to understand disease pathobiology.

## Materials and Methods

### Patient recruitment and Sample Collection

Patient samples were collected between 1 May 2020 and 9 June 2021 from patients seen at the Keck Hospital, Verdugo Hills, and Los Angeles (LA) County Hospital and stored in the University of Southern California (USC) COVID-19 Biospecimen Repository. At this time, no subjects were vaccinated nor were samples analyzed for SARS-CoV-2 variant. For this study, patients were assigned anonymized, coded IDs and were grouped according to the following cohort definitions: severe, indicating COVID-19 positive subjects who were admitted to the intensive care unit (ICU); moderate, indicating COVID-19 subjects who were hospitalized, but not admitted to the ICU; mild, indicating COVID-19 subjects who tested positive for SARS-CoV-2 but did not require hospitalization; and control, indicating subjects who tested negative for SARS-CoV-2 upon admission to the ICU for treatment of other severe diseases. Population demographics for these cohorts have been previously published [29]. Participants were predominantly Hispanic/Latinx (69%), reflecting the demographics of donors available from the source biorepository (57.4% Hispanic/Latinx, https://sc-ctsi.org/about/covid-19-biorepository).

### Proteomics

Plasma proteomics datasets have been previously published [29]. In brief, plasma samples were analyzed by Olink proximity extension assays (PEA) for quantification of 184 secreted markers. Olink’s Target 96 Inflammation and Target 96 Oncology II panels were chosen for the spread of proteins related to immune response and tissue remodeling. Of the 184 proteins in the panels, 6 were duplicates and 7 had NPX values under the protein-specific limit of detection (LOD) in >50% of samples in all cohorts, leaving 171 unique proteins for analysis. In total, 144 samples were analyzed. Samples were determined to fail quality control if internal incubation and detection controls deviated +/- 0.3 Normalized Protein eXpression (NPX) value from the median value across all samples. Four samples failed both panels and were excluded and eight samples failed the Oncology II panel and were only included in the analysis of the Inflammation panel.

### RNA Extraction & Quantitation

Total RNA was extracted from whole blood samples using the MagMAX for Stabilized Blood Tubes RNA Isolation Kit (Thermo Fisher Scientific, Waltham, MA), respectively, according to the manufacturer’s high-throughput protocol using the KingFisher Duo Prime Purification System (Thermo Fisher Scientific). RNA was eluted in 50 μl of MagMAX Elution Buffer, and yield was determined by quantitative real-time PCR using the TaqMan Fast Virus 1-Step Master Mix (Applied Biosystems, Foster City, CA) and TaqMan Gene Expression Assay, GUSB (Applied Biosystems), using Promyelocytic Leukemia (HL-60) Total RNA (Invitrogen) as the standard, according to the manufacturer’s recommendation. A concentration of 10 ng in 7 μL, or 1.43 ng/μL, was required as an adequate yield to proceed to cDNA synthesis. Total RNA was reverse transcribed using the SuperScript VILO cDNA Synthesis Kit (Invitrogen) according to the manufacturer’s specifications.

### Library Preparation and Next-Generation Sequencing

RNA libraries were prepared from reverse-transcribed cDNA samples on the Ion Chef System using the Ion AmpliSeq Kit for Chef DL8 and Oncomine Immune Response Research Assay (Thermo Fisher Scientific). RNA libraries were immediately used for sequencing. Magnetic bead purification and size-selection steps were performed using AgenCourt AMPure XP Beads (Beckman Coulter, Brea, CA) and DynaMag-PCR Magnet (Invitrogen). Sequencing of prepared RNA libraries was performed on the Ion Chef and Ion GeneStudio S5 Systems. RNA libraries were sequenced using the Ion 520 and Ion 530 Kit and Ion 530 Chips.

### NGS Analysis Pipeline and QC

Base calling, alignment, read filtering, and variant calling was performed on the IonTorrent Suite (v5.16). Reads smaller than 25 bases were removed. Thumbnail quality control reports produced by the Ion Torrent Suite were assessed for percentage ion sphere particle (ISP) loading and density, total reads and percentage usable reads, and read length. Runs were excluded if percentage ISP loading was below 70% overall or if ISP density was below 50% in any region of the chip, if percentage usable reads was below 70%, or if median read length was below 120bp. RNA libraries were aligned to the Immune Response (v3.1) reference library and analyzed using the ‘immuneResponseRNA’ plugin on the Ion Torrent Suite.

### Single-omics data analysis

Unpaired student’s t-test with p value ≤ 0.05 and adjusted p value (FDR) ≤ 0.1 were applied. Additionally, a fold-change cut-off was employed to obtain the differentially expressed features. Differentially expressed proteins (DEPs) were subjected to hierarchical clustering analysis, volcano plot, and Principal Component Analysis (PCA) using Olink Statistical Analysis Application (v1.0). Gene ontology and pathway enrichment analysis were retrieved from KEGG and Reactome using g:profiler [30]. Transcriptome Analysis Console (TAC, Applied Biosystems, v4.0.1.36) was used to perform one-way ANOVA comparison of gene expression levels from the RNA-seq data set. TAC was used to perform hierarchical clustering, generate a heatmap of gene expression levels, and generate volcano plots. Distances for hierarchical clustering were computed using the complete linkage method. Network analysis was performed using the STRING database (STRING Consortium, version 11.5). Protein-protein connections were assigned a combined “score” by evaluating probabilities of interaction derived from literature and database mining, then mapped according to these scores. The minimum required interaction score was set at the highest confidence (0.9) [31].

### Dataset Preparation for mixOmics analysis

Both RNA and proteomic datasets required data cleaning prior to model building. For RNA data from Ion Torrent, individual excel sheets containing log2-scaled, housekeeping normalized counts for up to 8 samples were imported into R. ‘Tidyverse’ functions were used to merge these tables into one ‘feature by sample’ matrix with redundant gene names corrected. For proteomic data, the Olink batch-normalized data in excel format was read into R and filtered to retain only patient’s plasma samples. Sample name errors were fixed at this stage. The resulting data tables for RNA and proteomics were written to CSV format for use in downstream modeling.

### Sparse PLS modeling of RNA-seq data alone

Our study included 65 matched samples with both RNA-seq and proteomic data available; the data had 398 features (transcriptomics) and 184 features (proteomics) as inputs to the model. Both datasets were normalized according to the default software protocols, as described previously. Sparse Partial Least Squares Regression Discriminant Analysis (sPLS-DA) models on the RNA-seq data alone were calculated with and without the covid-negative control samples included. An initial model with covid-negative samples (N=65) was created with feature selection disabled using ten components. A subsequent model with covid-negative samples removed (N=55) was subjected to performance testing with K-fold cross validation (K=5) and 50 repeats. The results showed the lowest error for five components (no feature selection). Feature selection tuning was then performed with K-fold cross validation (K=5) and 10 repeats, using the ‘Balanced Error Rate’ (BER) and ‘max.dist’ as the measure of performance. Feature tuning showed that just two components performed as well (BER < 15%) as 3 or more for certain values of “keepX” (see **Supplementary Figure S1**). The final sPLS-DA model on the RNA-seq data alone was constructed with just two components (to reduce risk of overfitting) and selected features of 40 and 50 on each component.

### MixOmics multi-omics data integration

The proteomic and transcriptomic datasets were integrated with ‘Data Integration Analysis for Biomarker Discovery using Latent components (DIABLO)’, a multiomics method that maximizes the correlation between pairs of pre-specified omics datasets using sparse PLS-Discriminant Analysis. Our study included 55 matched samples (as described above) as inputs to the model. Both datasets were normalized according to the default software protocols, as described previously. A 2x2 matrix was used as the design matrix to tune the model towards prioritizing sample classification performance versus maximizing feature correlations (values can range between 0 and 1):

|  | RNA | Proteomics |
| --- | --- | --- |
| RNA | **0** | **0.1** |
| Proteomics | **0.1** | **0** |

Model building was also explored with a range of off-diagonal values including 0.5 (balance of classification and correlation) and 0.75 (bias toward correlation); however, we did not observe a pronounced effect on the features selected or sample classification performance (data not shown). An initial DIABLO model was fit with ten components, the design matrix described above, and no feature selection for tuning and evaluation. Performance testing with K-fold cross validation (K=5) and 50 repeats showed that the overall balanced error rate (BER) decreased with each component until leveling out around 8 components (**Supplementary Figure S2A**). Thus, we carried eight components into tuning for feature selection with the ‘tune.block.splsda’ function (**Supplementary Figure S2B**). Tuning was performed with K-fold cross-validation (K=5) and ten repeats using ‘centroid distance’ measures. An optimal number of features was reported for each of 8 components across both blocks, with the BER approaching ∼0.1 for the optimal solutions. Performance evaluating the DIABLO model again after feature selection optimization showed that a minimum in the BER (∼0.1) was now reached at only four components. We used four components in constructing the final DIABLO model, retaining (20, 25, 25, 25) and (5, 7, 5, 5) features for each of four components in the RNA and proteomic data, respectively.

### Gene Ontology and Reactome Pathway Analysis

Gene ontology (GO term) analysis of the features selected by the DIABLO model as significantly correlated with and predictive of COVID severity for the RNA-seq (N_features=91) and proteomics (N_features=22) datasets was conducted using the Gene Ontology Resource Portal (geneontology.org; PANTHER v17.0, GO Database 2022-07-01). Fisher’s Exact test was used for overrepresentation analysis and an FDR correction (Benjamini and Hochberg) was applied. 103 out of 110 IDs were uniquely mapped (11 were multi-mapping). Pathway analysis of the same set of genes was performed with Reactome pathway browser (reactome.org; v3.7, database release 83).

### Study approval

The study was approved by the institutional review board (IRB) of the University of Southern California (USC): Protocol#: HS-20-00519.

## Results

In this study we examined a cohort of 70 patients which included four independent sub-groups. These comprised: 1) ‘COVID-positive-ICU’, our most severe response to SARS-CoV-2 infection with patients requiring treatment in the intensive care unit (ICU), 2) ‘COVID-positive-Inpatient’, our moderate group comprising of patients infected with SARS-CoV-2 requiring hospitalization, 3) ‘COVID-positive-outpatient’ samples, representing our mildest COVID-19 infections where patients tested positive for SARS-CoV-2 but required no hospitalization and 4) “Non-COVID-ICU’, a control group of ICU inpatients not infected with SARS-CoV-2. In this study we refer to these cohorts as severe, moderate, mild, and negative, respectively, for simplicity. An overview of the experimental design is presented in **Fig. 1** and demographic information of the study cohorts can be found in [29].

**Figure 1.**
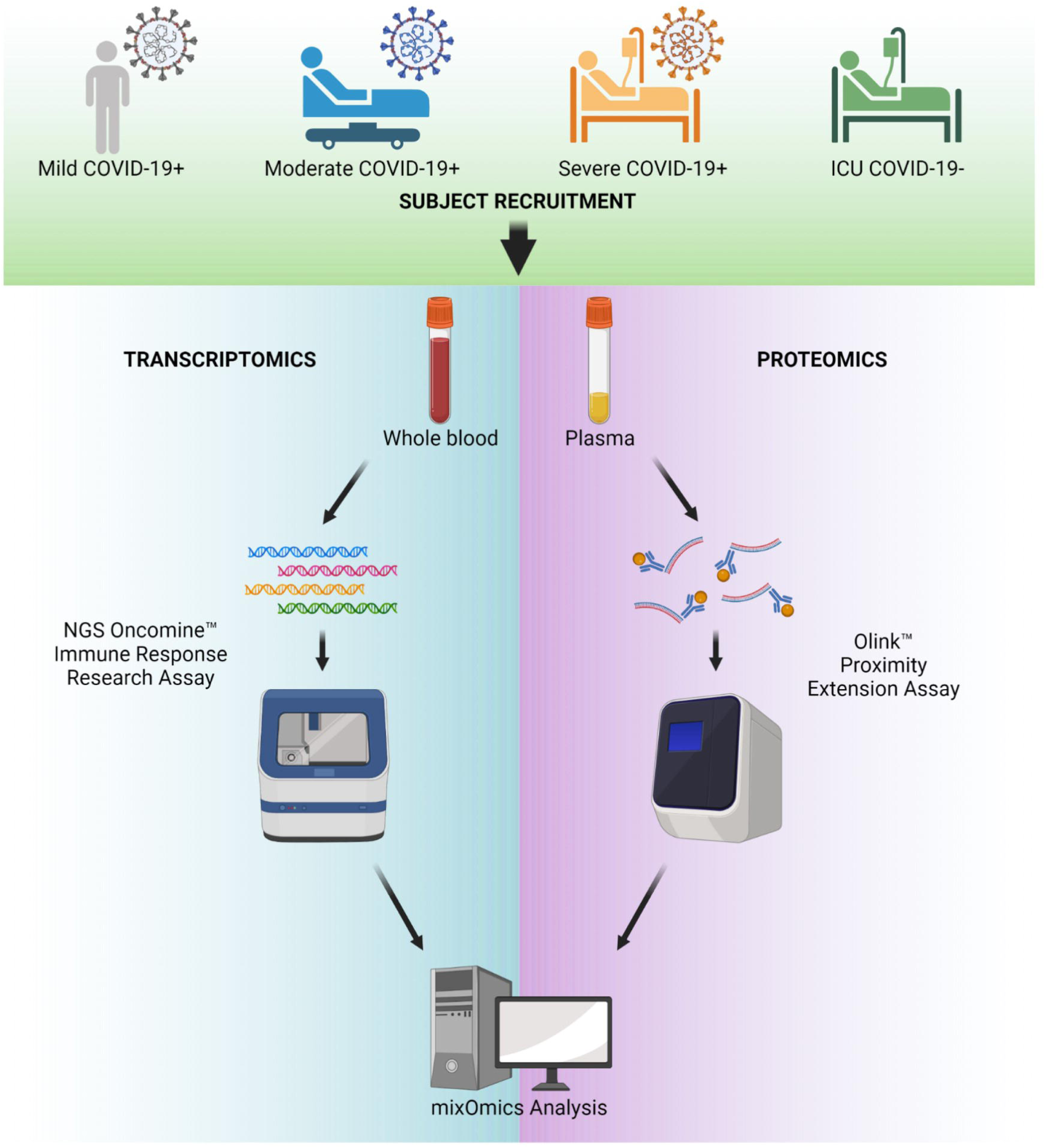
COVID-19 multiomic study design. Schematic diagram of experimental design for the study.

### Transcriptomic analysis of COVID-19 severity

To evaluate transcriptomic changes, we used a targeted next generation sequencing (NGS) Oncomine Immune Response Research Assay to quantify the expression of 395 genes specifically associated with inflammatory signaling and immune-oncology research. This automated workflow allows for reproducibility across samples and requires a low RNA input allowing for evaluation of repeat blood samples from COVID-19 patients. Unsupervised clustering of gene expression shown in the heatmap in **Supplementary Figure S3A** represents the gene expression from the first sample collected from all subjects in the severe, moderate, and mild COVID-19 cohorts and highlights three major clusters of gene expression representing mild/moderate (purple/red), severe/moderate (blue/red) and severe COVID-19 (blue). From the targeted Oncomine-Immune panel of 395 genes evaluated in this study there are 181 significant differentially expressed genes (DEG) comparing severe to mild, 68 DEG comparing severe from moderate, and 104 DEG comparing moderate to mild COVID-19 cohorts (**Supplementary Figure S3B-C and Supplementary Database S1**). Among the DEG there are 67 genes that separate severe COVID-19 from both moderate and mild cases (**Supplementary Database S1 and Supplementary Figure S3C**). Comparison of transcript levels across the top 25 most DEG between severe and all cohorts (**Fig. 2A**) shows 4 clusters of gene expression: 1) genes with low expression in mild and with decreased expression with severity, 2) genes highly expressed in mild and with decreased expression with severity, 3) genes expressed in mild and highly decreased with severity and 4) genes expressed in mild and greatly increased with severity. STRING (Search Tool for the Retrieval of Interacting Genes/Proteins) evaluation of the physical and functional protein-protein interaction network of the top 25 most significant DEG between severe and all other cohorts (**Supplementary Database S1**) highlights two major clusters of predicted interactions centering around mTOR-FOXO1 signaling and PTPRC signaling. The colored nodes are proteins in our dataset and the white nodes are proteins with predicted interactions in these networks (minimum interaction score of 0.7) (**Fig. 2B**). The bar charts provide examples of severity-dependent DEG in COVID 19 (**Fig. 2C-E**). ARG1 and S100A9 both increase in expression with severity, with similar trends in both male and female subjects (**Fig. 2C**). HLA-B and IFNA17 are genes that are increased with severity, but only in distinct clusters of the cohort. Interesting HLA-B expression is differentially associated with severity in male and female subjects with significantly elevated expression associated with moderate cases specifically in females and not in males (**Fig. 2D**). CLEC4C is an example of a DEG that is decreased with severity with exceptionally low expression in severe COVID-19 cases (**Fig. 2D**).

**Figure 2.**
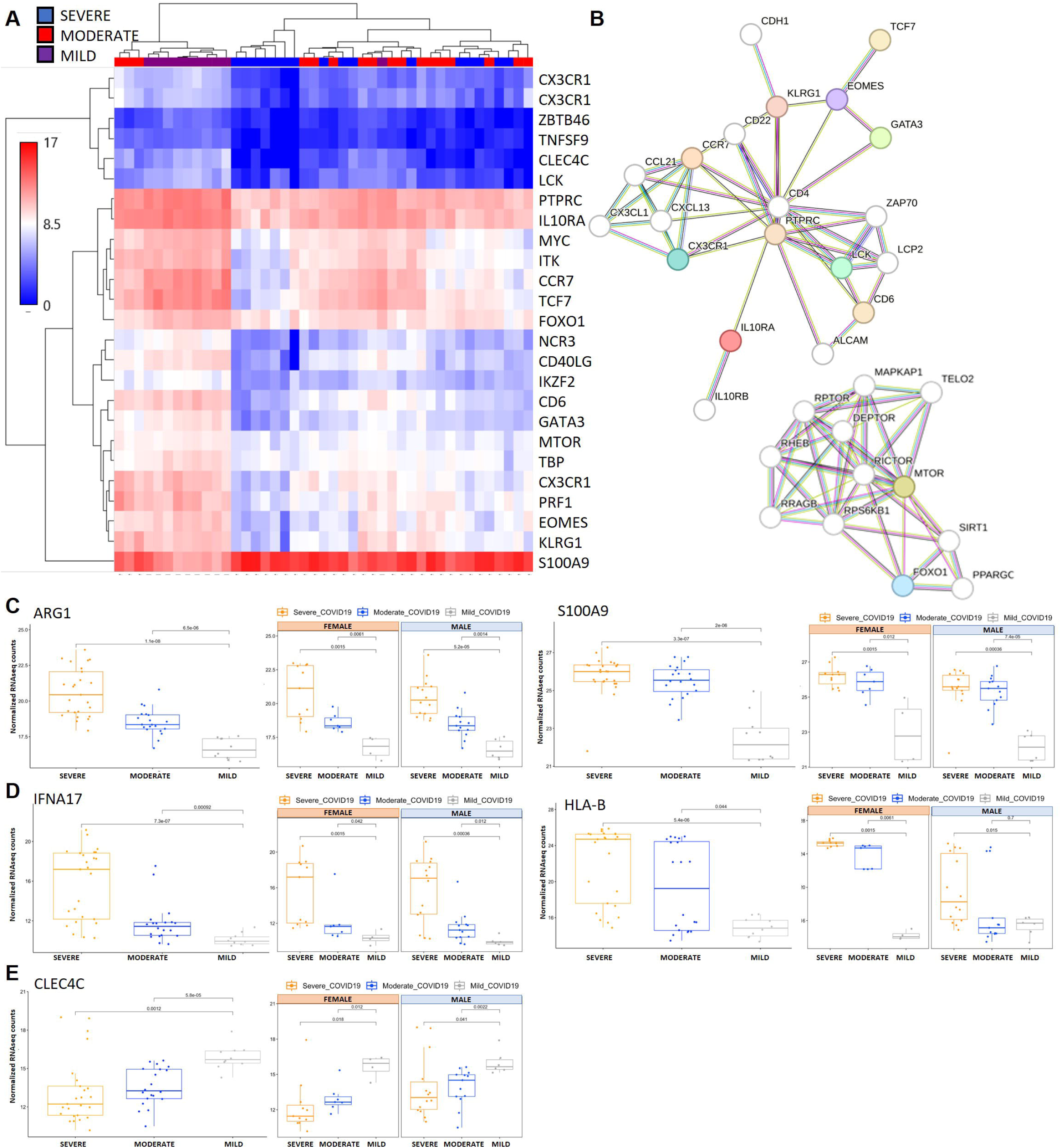
RNAseq analysis comparing all COVID-19 cohorts. **(A)** Heatmap showing unsupervised clustering of the top 25 most significant DEG between severe and all other COVID-19 study cohorts. Relative expression is on the scale of 0 (blue) to 17 (red) for COVID-19 cohorts severe (blue), moderate (red) and mild (purple). **(B)** STRING analysis showing predicted protein-protein interactions between the top 25 DEG from Severe COVID-19 compared to all other cohorts. Colored nodes represent query proteins, white nodes represent second shell of interactions. Known interactions are shown from curated databases (teal lines) or experimentally determined (pink lines). Predicted interactions shown are based on gene neighborhood (green lines), gene fusions (red lines), gene co-occurrence (blue lines). **(C-E)** Bar charts comparing Log2 fold change in average transcript level across Day 1 samples from all subjects in severe, moderate, and mild COVID-19 cohorts. Examples of significantly DEG include S100A9 and ARG1, consistently elevated with severity of COVID-19 (**C**), CLEC4C an KRT5, consistently decreased with severity of COVID-19 (**D**) and HA-B and IFNA17, elevated with severity of COVID-19 in a portion of the subjects within the severity category (**E**).

### Single ‘omics’ modeling with sparse PLS discriminant analysis

Next, we performed a sparse partial least squares model with discriminant analysis (sPLS-DA) that separated severe, moderate, and mild COVID-19 (**Fig. 3A**), but not the ‘COVID-19-positive-inpatient’ (moderate) from ‘COVID-19-negative-inpatient’ (negative) (**Fig. 3A**). We removed the COVID-19-negative inpatient samples since they are not directly relevant to the investigation of biomarkers of COVID-19 severity; this had no impact on the ability of the model to predict COVID-19 status on RNA sequencing (RNA-seq) expression data (**Fig. 3B**). Classification receiver operating characteristic (ROC) curves for the RNA-seq sPLS-DA model, with the COVID-19 negative samples removed, illustrates the diagnostic ability of the model with values of 0.98 for severe versus all other samples, 0.97 for moderate vs all other samples, and 1.0 for mild vs all other samples (**Fig. 3C**). The sPLS-DA model of the RNAseq data alone demonstrates nearly perfect clustering and prediction of COVID-19 severity based on just 2 components, comprised of 40 and 50 features in the data, respectively (**Fig. 3**).

**Figure 3.**
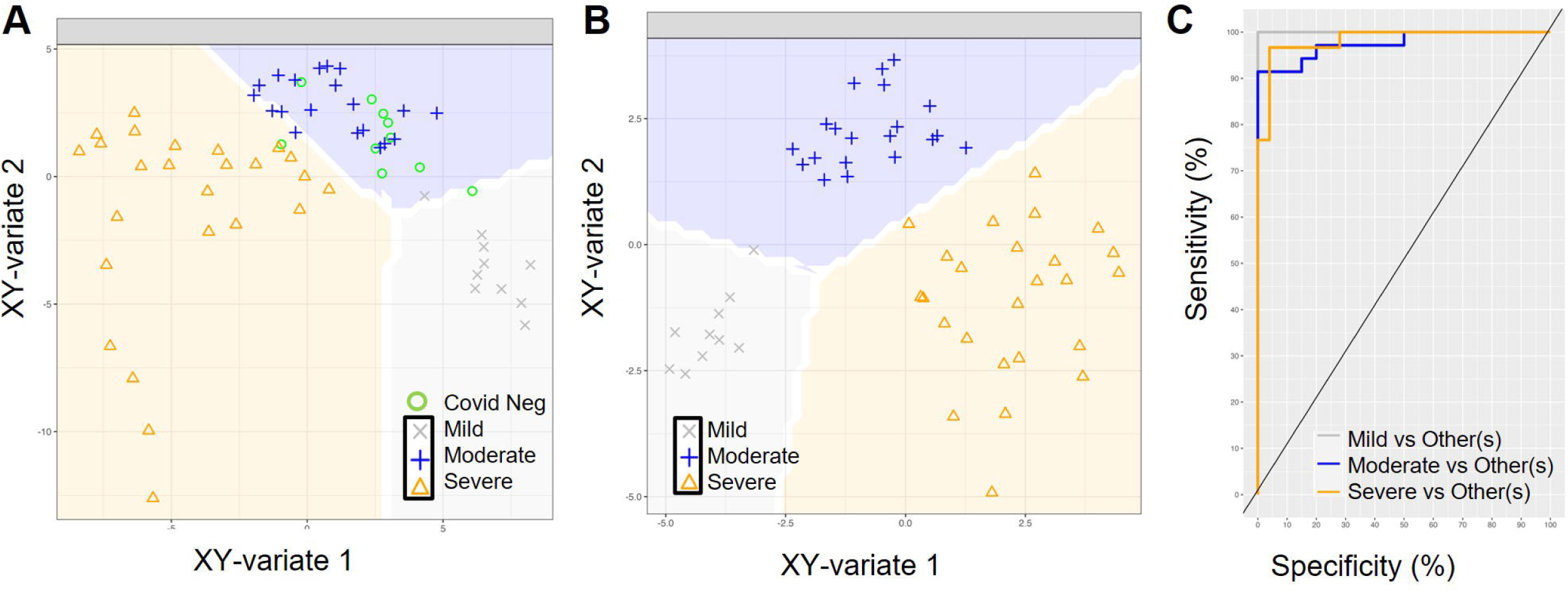
sPLS Discriminant Analysis for classification of COVID-19 severity. RNA-seq single dataset sPLS-DA component plots with decision background with **(A)** and without **(B)** COVID negative samples. Samples are projected onto their XY-Variate latent spaces using only the RNA-seq data and are colored by COVID-19 status. The prediction background generated by the model is plotted behind the samples, showing decision boundaries for classifying new samples. **(C)** ROC analysis of the model in (B) showing very high AUCs for each sample category.

### ‘Omics’ integrative modeling with DIABLO

As transcriptomics and proteomics are interrelated layers of the overall system that determines a cells response to SARS-CoV-2 infection, we performed a multivariate analysis (described in the Methods) to integrate the changes observed in the proteomics data (described previously in [29]) and the transcriptomic data described above. We applied the Data Integration Analysis for Biomarker discovery using Latent variable approaches for ‘Omics studies method (DIABLO) of the MixOmics package which applies the sPLS-DA model in the context of two or more related datasets on the same set of samples. Modeling the COVID-19 positive samples demonstrated a clear separation of severe, moderate, and mild COVID-19 patient samples when observing latent variables (“Components”) 1 and 2 using a weighted average of both blocks (**Fig. 4A**). Introspection of the features selected by the DIABLO model enables the identification of key molecular drivers from our dataset in the context of COVID-19 severity. The top 117 features selected by the DIABLO model are reported in **Supplementary Table S1.** We note that the first component (‘variate 1’) primarily separates the mild samples from the rest, while the second component partially (‘variate 2’) separates the moderate from the severe. The top features (genes and/or proteins) that contribute to each of the ‘blocks’ of the first and second component (of four total components) are shown in **Fig. 4B-C**. The top features contributing to the classification performance of the final DIABLO model along the first component in the RNA dataset include genes IL2RB, ARG1, CD4, IL10RA, CA4, MYC, CD6, CSF1R, TCF7, ZAP70, ITK, S100A9, CD8B, SIT1, and FCGR1A and SYND1, EN-RAGE, and WFDC-2 in the proteomic dataset. This group of genes and proteins forms a highly enriched protein-protein network (STRING analysis PPI enrichment p-value 1^-16) with functional enrichments in IL-15 signaling (GO: Biological Processes), T-cell receptor complex (GO: Cellular components) and KEGG pathways for primary immune deficiency and Th1 and Th2 cell differentiation (**Supplemental Database S2**). The second component is comprised of top-weighted genes IRS1, CCR4, TFRC, CD79A, IGSF6, SELL, MIF, IL15, CD19, CXCR2, IFITM1, and AIF1 in the RNA data and GZMB, CD70, SPARC, CD5, and LYPD3 in the proteomic dataset (**Figure 4D-E**). This group of genes and proteins forms a highly enriched protein-protein network (STRING analysis PPI enrichment p-value 2.75^-14) with functional enrichments in negative regulation of myeloid cell apoptosis and neutrophil activation (GO: Biological Processes), plasma membrane and cell surface (GO: Cellular components) (**Supplemental Database S3**).

**Figure 4:**
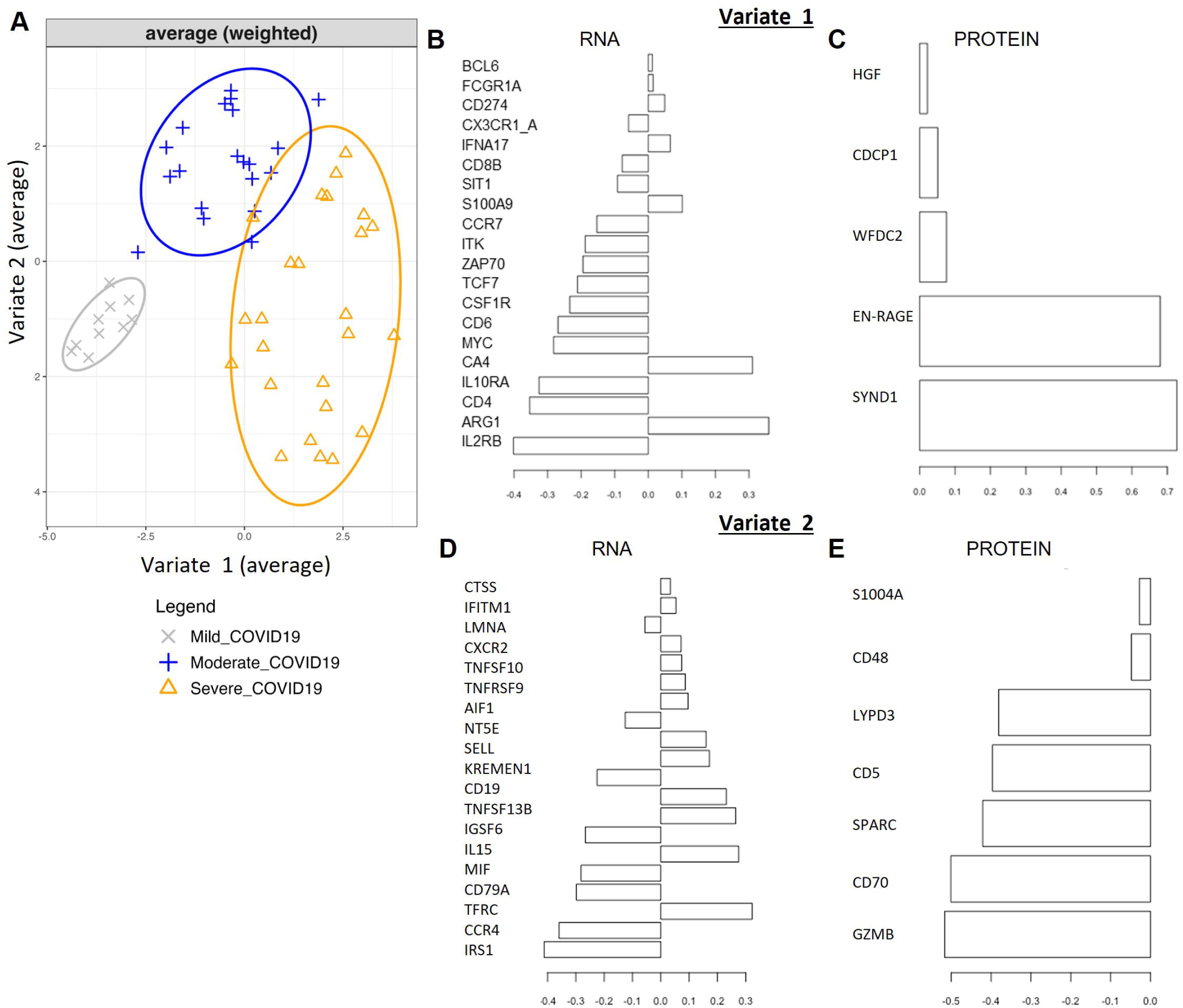
MixOmics Integration of the RNA and Protein datasets using DIABLO. **(A)** PCA plot of component 1 (Variate 1) and component 2 (Variate 2) of 4 components used to define sample clusters. **(B-E)** Top genes **(B and D)** and proteins **(C and E)** defining the clustering of the samples based on component 1 (B-C) and component 2 **(D-E).** The X-axis represents the “loading” on each feature: a measure of how important it is to the trained model. This is a vector of the weight of each original variable’s contribution to the corresponding “latent” variable (Variate 1, Variate 2, etc.).

Circos style plots in **Figure 5A** show how selected RNA and proteomic features, having the largest loadings and cross-block correlations, in Component 1 are positively and negatively correlated to each other. The “Clustered Image Map” (CIM) heatmap shown in **Figure 5B** highlights the correlation strength between a given pair of features represented by the differences in “cell” color. The CIM visualizes the correlation structure extracted from both the RNA and proteomic datasets. Blocks homogeneous in color depict subsets of features from each dataset that are correlated and is suggestive of a potential causal relationship. Visualizing this data as a network plot provides additional context to the correlation between features (**Fig. 5C**). The strength of a positive (red) or negative (green) correlation between core features defining the dataset clustering is shown in the lines connecting proteins (green) and RNA (blue). The core of these correlations center around connections with Syndecan-1, EN-RAGE, WFDC2, HGF and CDCP1. Unbiased, hierarchical clustering of gene and protein expression data based on genes used to construct the two components of the sPLS-DA model shows almost perfect separation of severe and mild COVID-19 samples (**Figure 6**), highlighting the strength of this model to predict signatures associated with COVID-19 severity.

**Figure 5.**
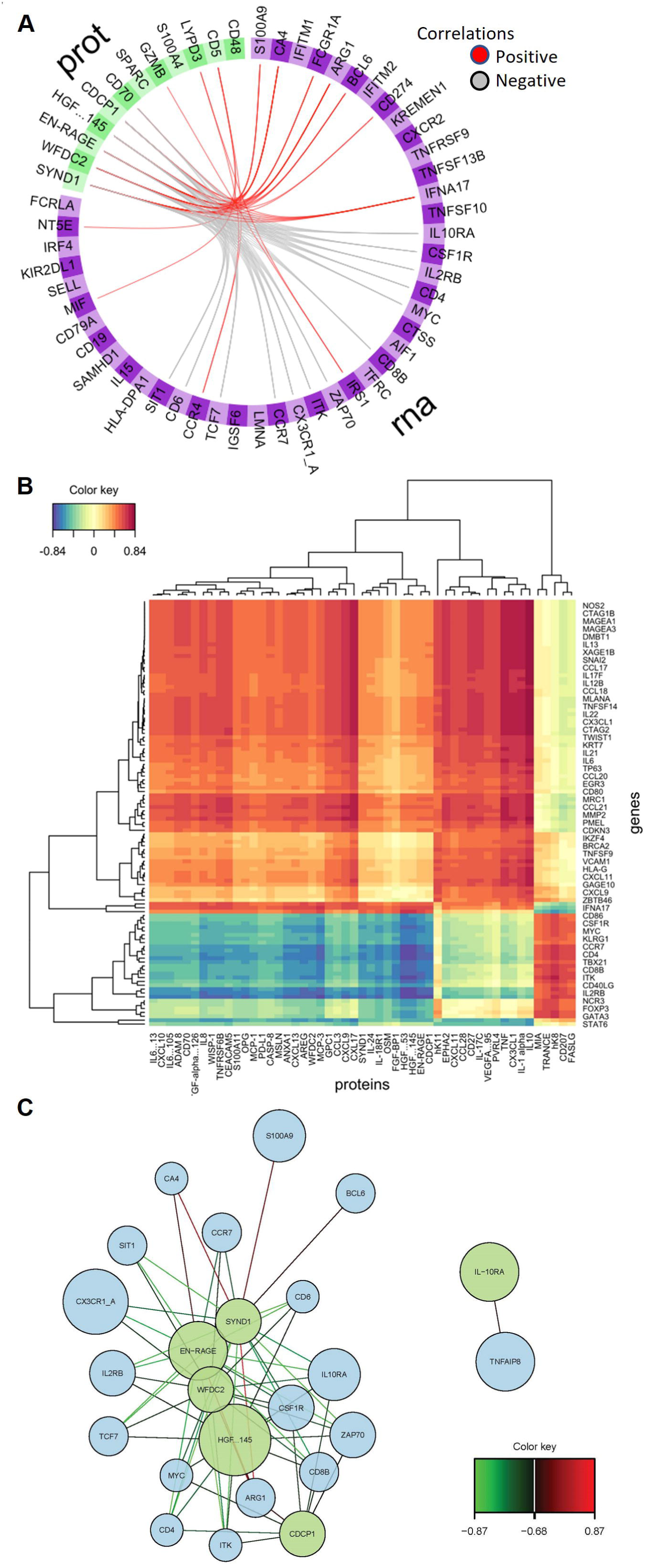
Positively and negatively correlated features of the datasets. **(A)** Circos plot showing highly positively and negatively correlated features between the RNA and protein datasets (with correlation cutoff of 0.65). The two different datasets are segmented and colored across the circle with each subsection representing a specific feature. The lines within the circle represent positive or negative correlations between linked variables. **(B)** Clustered expression heatmap of the highly correlated features in the DIABLO sPLS-DA model. Both features (Y-axis) and samples (X-axis) are clustered in an unsupervised manner. **(C)** Network plot of highly correlated variables where the connections represent correlations in the data (red is positive correlation and green is negative correlation). Genes are found in the blue circles and proteins in the green circles.

**Figure 6:**
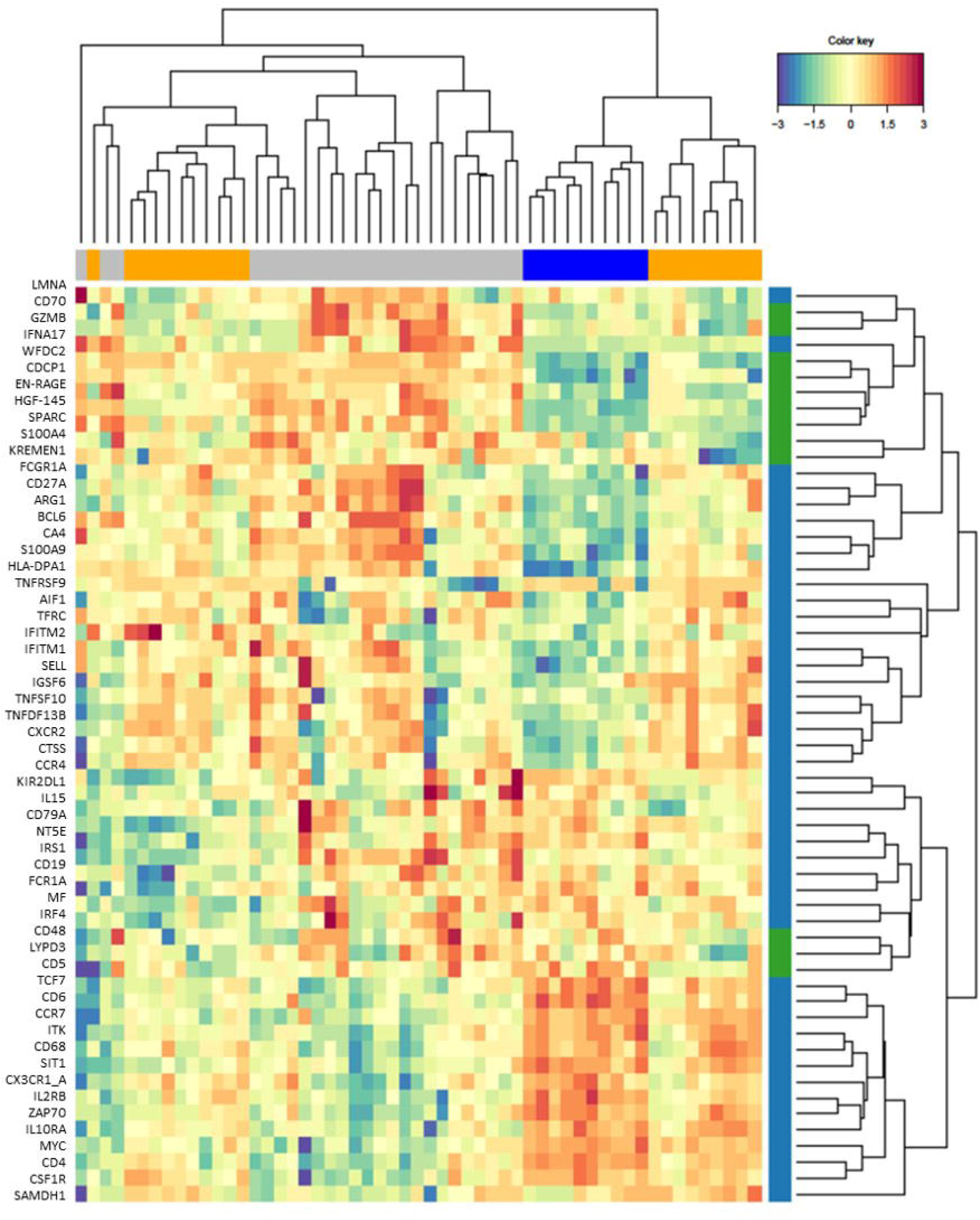
Highly correlated dataset features that maximizing connections to outcomes. Heatmap generated based on analysis of all features and removing non-informative features to maximally connect highly correlated variables to outcome. Proteins (green) and genes (blue) are depicted on the y-axis and cohorts, grey (mild), blue (moderate) and orange (severe) are shown on the x-axis. The scale represents a relative expression from -3 (blue) to 3 (red).

### GO term and pathway analysis of selected biomarkers of COVID-19 severity

As described in *Methods*, the final DIABLO model selected 95 and 22 features from the RNA-seq and proteomics datasets, respectively, after model tuning. These ∼117 features were used as inputs to Gene Ontology (GO) term analysis (**Table 1**) and Reactome pathway analysis (**Table 2**) to identify ontologies and pathways that are enriched in our COVID severity biomarkers. Because we are starting from a panel of genes already selected for oncology and inflammation, we expect to see enrichment for general or high-level terms and pathways. Therefore, we only considered GO terms and pathways an FDR-corrected p-value < 0.001 and fold enrichment >10 (GO terms) as potentially significant. GO term analysis showed an enrichment in terms related to “immune system process,” as expected. However, terms with the largest fold enrichments (>10) as well as significant FDR p-values were related to regulation of “lymphocyte activation”, “T-cell activation”, “leukocyte activation”, and “leukocyte proliferation.” Among the enriched pathways from ReactomeDB analysis, only four had FDR-corrected p-values less than 0.001 (“Interleukin-4 and Interleukin-13 signaling”, “Cytokine Signaling”, “Immune System” and “Interleukin-10 signaling”).

**Table 1.** Top 25 GO terms enriched among DIABLO-selected features.

| GO biological process complete | Homo sapiens - REFLIST (20589) | Upload # (110) | Upload (expected) | Upload (over/under) | Upload (fold Enriched) | Upload (raw P-value) | Upload (FDR) |
| --- | --- | --- | --- | --- | --- | --- | --- |
| immune system process (GO:0002376) | 2429 | 73 | 12.98 | + | 5.63 | 1.08E-40 | 1.69E-36 |
| regulation of immune system process (GO:0002682) | 1520 | 56 | 8.12 | + | 6.9 | 1.57E-33 | 1.23E-29 |
| immune response (GO:0006955) | 1621 | 55 | 8.66 | + | 6.35 | 4.74E-31 | 2.48E-27 |
| positive regulation of immune system process (GO:0002684) | 967 | 43 | 5.17 | + | 8.32 | 5.24E-28 | 2.05E-24 |
| regulation of lymphocyte activation (GO:0051249) | 591 | 34 | 3.16 | + | 10.77 | 2.86E -25 | 8.98E-22 |
| regulation of T cell activation (GO:0050863) | 377 | 29 | 2.01 | + | 14.4 | 8.19E-25 | 2.14E-21 |
| regulation of leukocyte activation (GO:0002694) | 684 | 35 | 3.65 | + | 9.58 | 2.00E-24 | 4.47E-21 |
| regulation of cell activation (GO:0050865) | 741 | 35 | 3.96 | + | 8.84 | 2.49E-23 | 4.89E-20 |
| regulation of immune response (GO:0050776) | 935 | 37 | 5 | + | 7.41 | 3.54E-22 | 6.16E-19 |
| leukocyte activation (GO:0045321) | 581 | 31 | 3.1 | + | 9.99 | 4.52E-22 | 7.09E-19 |
| response to stimulus (GO:0050896) | 8209 | 93 | 43.86 | + | 2.12 | 7.35E-22 | 1.05E-18 |
| response to organic substance (GO:0010033) | 2704 | 55 | 14.45 | + | 3.81 | 2.32E-20 | 3.03E-17 |
| cell activation (GO:0001775) | 700 | 31 | 3.74 | + | 8.29 | 8.05E-20 | 8.41E-17 |
| regulation of cell population proliferation (GO:0042127) | 1674 | 44 | 8.94 | + | 4.92 | 7.79E-20 | 8.73E-17 |
| cellular response to stimulus (GO:0051716) | 6569 | 82 | 35.1 | + | 2.34 | 7.34E-20 | 8.86E-17 |
| cell surface receptor signaling pathway (GO:0007166) | 2174 | 49 | 11.61 | + | 4.22 | 1.38E-19 | 1.35E-16 |
| regulation of leukocyte proliferation (GO:0070663) | 271 | 22 | 1.45 | + | 15.19 | 2.36E-19 | 2.18E-16 |
| defense response (GO:0006952) | 1478 | 41 | 7.9 | + | 5.19 | 3.50E-19 | 3.05E-16 |
| regulation of leukocyte cell-cell adhesion (GO:1903037) | 369 | 24 | 1.97 | + | 12.17 | 5.69E-19 | 4.46E-16 |
| lymphocyte activation (GO:0046649) | 465 | 26 | 2.48 | + | 10.47 | 5.54E-19 | 4.57E-16 |
| signal transduction (GO:0007165) | 4887 | 70 | 26.11 | + | 2.68 | 9.53E-19 | 7.12E-16 |
| positive regulation of T cell activation (GO:0050870) | 253 | 21 | 1.35 | + | 15.54 | 1.10E-18 | 7.84E-16 |
| regulation of response to stimulus (GO:0048583) | 4034 | 63 | 21.55 | + | 2.92 | 4.41E-18 | 3.00E-15 |
| positive regulation of leukocyte cell-cell adhesion (GO:1903039) | 276 | 21 | 1.47 | + | 14.24 | 5.78E-18 | 3.78E-15 |
| positive regulation of leukocyte proliferation (GO:007066) | 168 | 18 | 0.9 | + | 20.05 | 6.15E-18 | 3.86E-15 |

**Table 2.** Reactome Pathway Enrichment for DIABLO-selected features.

| Pathway identifier | Pathway name | #Entities found | #Entities total | #Interactors found | #Interactors total | Entities ratio | Entities pValue | Entities FDR | #Reaction found | #Reaction total |
| --- | --- | --- | --- | --- | --- | --- | --- | --- | --- | --- |
| R-HSA-6785807 | Interleukin-4 and Interleukin-13 signaling | 20 | 211 | 3 | 162 | 0.013845 | 1.37E-10 | 1.76E-07 | 9 | 47 |
| R-HSA-1280215 | Cytokine Signaling in Immune system | 65 | 1115 | 50 | 2999 | 0.073162 | 4.59E-08 | 2.95E-05 | 290 | 740 |
| R-HSA-168256 | Immune System | 88 | 2703 | 62 | 4209 | 0.177362 | 3.38E-07 | 1.44E-04 | 530 | 1659 |
| R-HSA-6783783 | Interleukin-10 signaling | 11 | 86 | 2 | 93 | 0.005643 | 6.72E-07 | 2.16E-04 | 13 | 15 |
| R-HSA-380108 | Chemokine receptors bind chemokines | 8 | 57 | 2 | 70 | 0.003740 | 4.74E-06 | 0.001212 | 12 | 19 |
| R-HSA-449147 | Signaling by Interleukins | 43 | 658 | 35 | 2161 | 0.043176 | 5.08E-05 | 0.010863 | 187 | 505 |
| R-HSA-389948 | PD-1 signaling | 5 | 45 | 1 | 4 | 0.002953 | 6.68E-05 | 0.012223 | 4 | 5 |
| R-HSA-202430 | Translocation of ZAP-70 to Immunological synapse | 5 | 42 | 3 | 14 | 0.002756 | 1.18E-04 | 0.018894 | 4 | 4 |
| R-HSA-9012546 | Interleukin-18 signaling | 3 | 11 | 1 | 5 | 7.22E-04 | 2.93E-04 | 0.041670 | 4 | 4 |

## Discussion

By applying MixOmics to combine approaches to interrogate high dimensional datasets we have been able to interrogate the molecular basis of COVID-19 disease severity more comprehensively. This integrative approach contrasts with most existing studies which only focus on a single ‘omics approach which likely masks valuable information. Our data highlights the power of combined analysis of independent transcriptomic and proteomic data sets taken from the same subjects across different disease severity cohorts to elucidate complex biological mechanism leading to severe COVID-19.

Integrative modeling discovered correlations between features previously found to be significantly differentially expressed and associated with COVID-19 severity in our proteomic data alone [29]. Application of DIABLO allowed us to identify key ‘omics variables from our transcriptomic and proteomic datasets and was able to discriminate between COVID-19 severity cohorts (**Fig. 6**). Importantly, the features that our model used to drive clustering of the datasets are consistent with data from other published ‘omics data analysis. Examples include elevated protein expression with severity of COVID-19 for proteins including: 1) Syndecan-1 (SYND1) [32–34], 2) S100 calcium binding protein A12 (EN-RAGE or S100A12) [35–37], 3) Hepatocyte Growth Factor (HGF) [38–40] and 4) CUB domain containing protein 1 (CDCP1) [41]. Examples for elevated RNA transcript expression with severity of COVID-19 for genes included 1) IFNA17 [42, 43], 2) ARG1 [44, 45]. Interestingly IFNA17 and HLA-B are two examples of genes where expression is associated with severity, but only in a sub-population of patients. HLA-B is known to exhibit significant genetic diversity among individuals and will influence one’s ability to recognize and respond to viral infection by COVID-19 [46, 47]. IFNA17 is an interferon that is a critical part of the innate immune response in viral infection. In support of this data, IFNA17 was discovered to be differentially expressed in a study evaluating interferon stimulated gene profiles of post-mortem lung tissues from severe cased of COVID-19 [42]. While its expression remains an active area of research in COVID-19 it is likely that its overexpression may lead to hyperinflammation in severe COVID-19 [48, 49].

While our study was designed to interrogate high-dimensional datasets where the patient sample may be limited, the small number of subjects in our study can also be viewed as a limitation. Furthermore, repeat samples were only obtained from inpatients in the USC COVID-19 Biorepository leading to an unbalanced design of the study cohorts. The study designed to investigate targeted panels of genes and proteins rather than taking a whole transcriptome and proteome approach. While this limits the scope of the target signatures associated with COVID-19 severity it allowed for the development of a multivariate integrative classification method that can predict signatures associated with COVID-19 severity that can be applied to integrate larger transcriptomic and proteomic datasets. The GO terms discovered through DIABLO highlighted a link between interleukin 4 and Interleukin 13 signaling and COVID-19 severity. Il-13 was recently discovered to be a core driver of COVID-19 severity; patients prescribed Dupliumab, an antibody that blocks IL-13 and Il-4 has significantly less severe disease. This observation was backed up by data in murine COVID-19 models [50]. Indeed, IL-13 signaling has been linked to the regulation of hyaluronic acid and the persistence of post COVID-19 conditions [50, 51]. Similarly, PD-1 and the PD-L1 axis has also been connected clinically to severity of COVID-19 [52–54]. PD-1 (CD279) is known to be involved in the maintenance of immune tolerance and several studies have now reported that regulation of the PD-1/PD-L1 axis is critical in the regulation of a variety of infectious diseases [55, 56]. While acutely a reduction in infection-associated inflammation and inflammation-mediated tissue damage may be noted, chronic activation can drive immune exhaustion and be associated with increased severity of infectious diseases, such as SARS-CoV-2.

These examples and the analysis presented in this study clearly demonstrates the capacity for MixOmics to discover correlations between the features of independent datasets and generate biomarker signatures specific to disease status. Wider application of this approach to published datasets should substantially enhance our ability to identify specific biomarkers predictive of COVID-19 disease severity and assist in understanding the biomolecular pathways defining the phenotype pathogenesis.

## Ethics approval and consent to participate

Patient samples were collected between 1 May 2020 and 9 June 2021 from patients seen at the Keck Hospital, Verdugo Hills, and Los Angeles (LA) County Hospital and stored in the University of Southern California (USC) COVID-19 Biospecimen Repository. The study was approved by the institutional review board (IRB) of the University of Southern California (USC): Protocol#: HS-20-00519 to access the samples in the USC COVID-19 Biospecimen Repository.

## Consent for publication

All authors have consented agreement for the publication of this study

## Availability of data and material

The proteomics dataset has been previously published [29] and the normalized protein expression (NPX) data provided for all samples by Olink is available at https://figshare.com/s/d136a74ef05c3dfa3a21. The Oncomine Immune Response Transcriptomic processed datasets and code to reproduce the DIABLO mixOmics analysis from data munging to model building can be found at the following link (https://github.com/mchimenti/covid_multiomics_mar2022).

## Author Contributions

N.C.H. conceived and performed the experiments, analyzed the data, and wrote the manuscript. M.C. developed and performed the MixOmics analysis and wrote the manuscript. A.L.R. conceived the study, conceived, and directed experiments, interpreted experimental data, and wrote the manuscript.

## Supporting information

Supplemental Database S3

Supplemental Database S2

Supplemental Database S1

## Acknowledgements

This research was made possible by the USC COVID-19 Biospecimen Repository, which provided patient plasma samples. We would like to thank Ben Darbro, University of Iowa, for the insightful discussions.

