## Supplemental Information for "MixOmics Integration of Biological Datasets Identifies Highly Correlated Key Variables of COVID-19 severity"

**SUPPLEMENRATY INFORMATION**

**Title**

**Affiliations**

Amy L. Ryan, PhD

Associate Professor: Anatomy and Cell Biology

Associate Director: Center for Gene Therapy

BSB, 1-400 Core

University of Iowa

51 Newton Road

Iowa City, Iowa 52241

**Conflict of Interest Statement**

The authors have declared that no conflict-of-interest exists.

**Supplementary Figures**

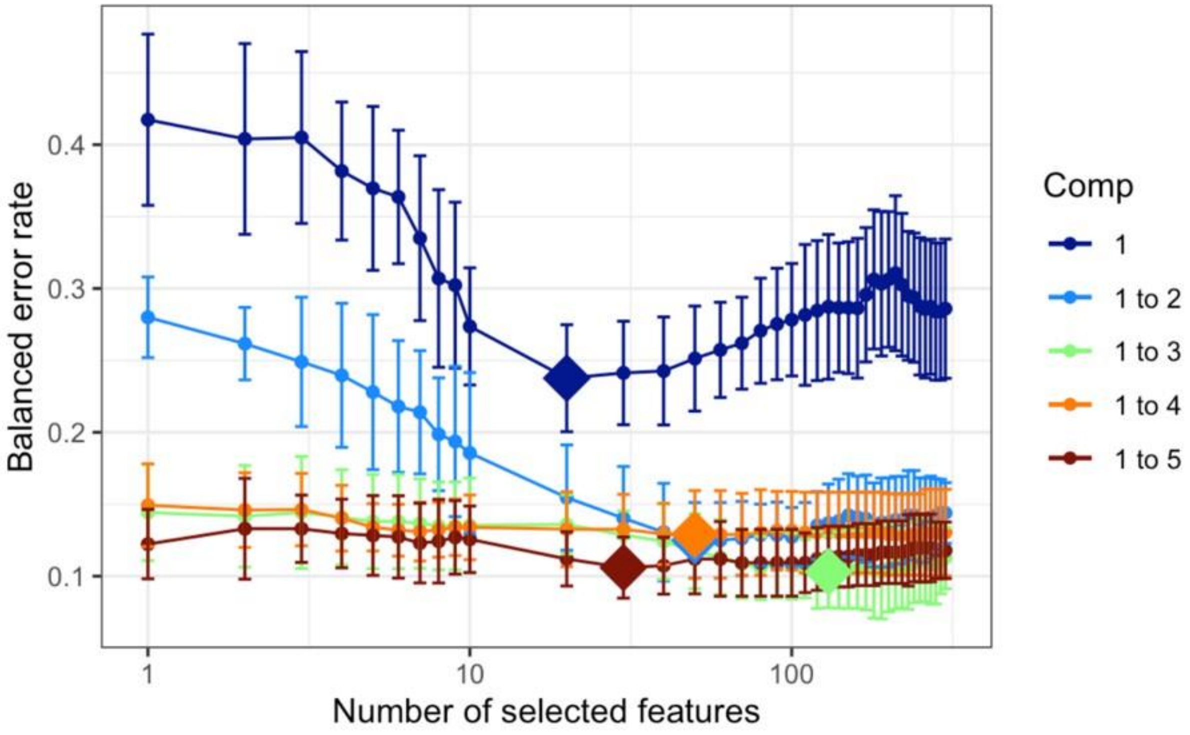

**Supplementary Figure S1. sPLA-DA feature selection tuning of the transcriptomics dataset.** 5-fold cross-validation of the sPLS-DA RNA-seq model with 10 repeats using a “balanced error rate (BER).” Plotting the BER as a function of the number of features selected showed that two components (light blue) performed nearly as well as 3-5 components for certain values of the “keepX” parameter.

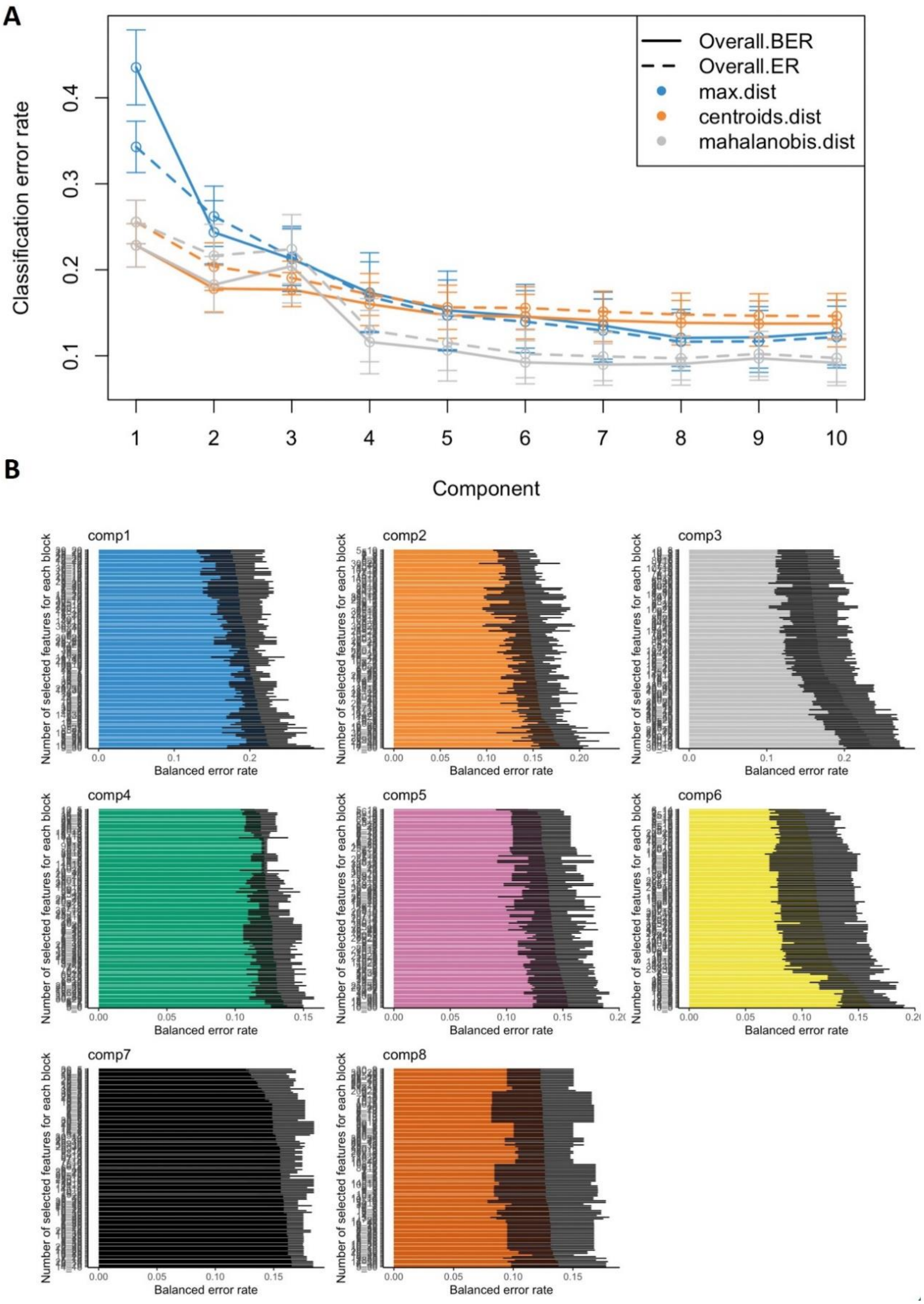

**Supplementary Figure S2. DIABLO model performance feature selection. A**) Performance testing with K-fold cross validation (K=5) and 50 repeats showed that the overall balanced error rate (BER) **decreased with** each component until leveling out around 8 components. **B**) Feature selection tuning on 8 components of the DIABLO model.

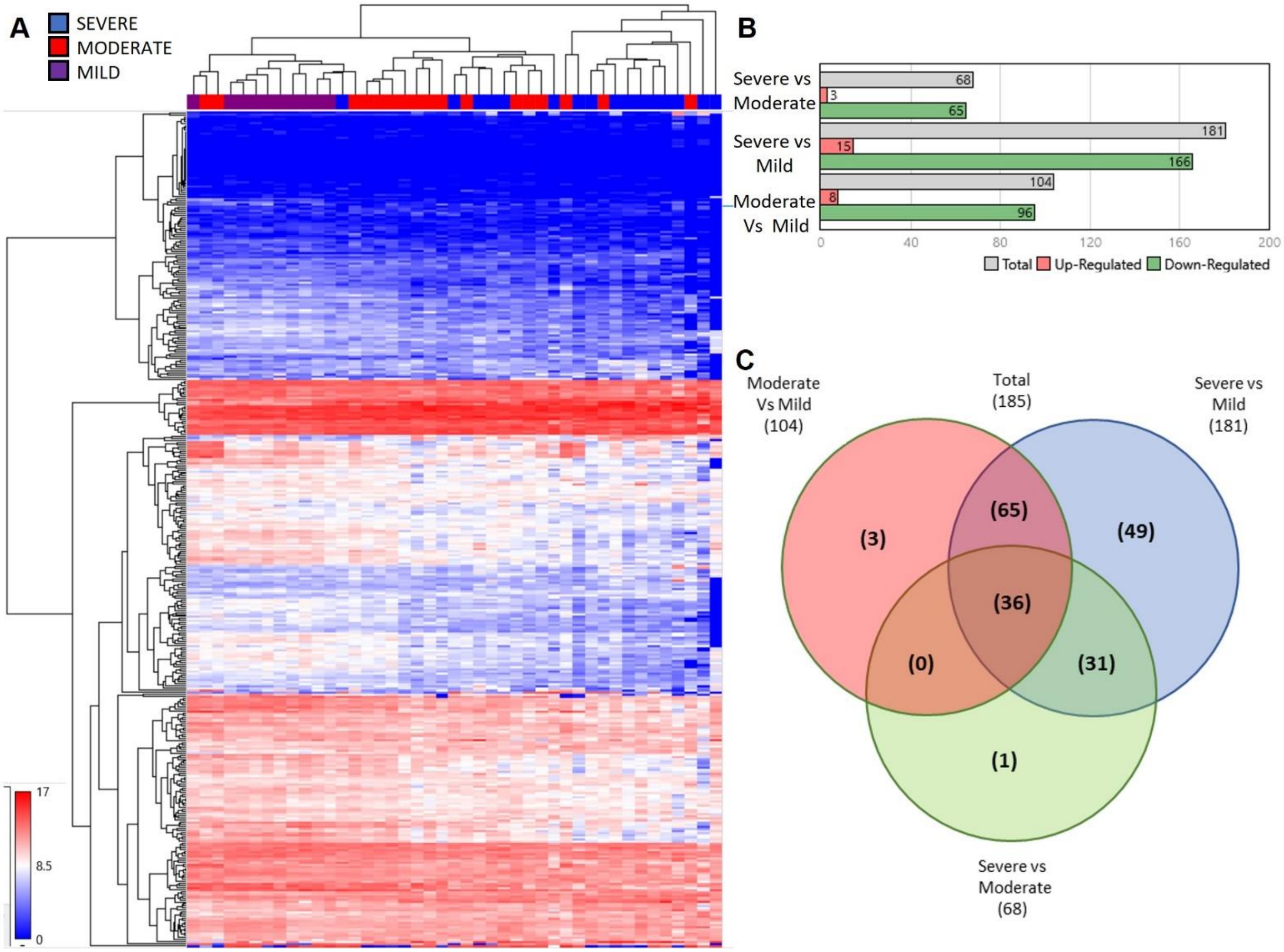

**Supplementary Figure S3. DEG between COVID-19 cohorts. A)** Unsupervised clustering of gene expression in a heatmap with cohorts identified as severe (blue) moderate (red) and mild (purple). **B)** Table of DEG between the paired cohort comparisons identified. Total DEG (grey), upregulated DEG (red) and down regulated DEG (green). **C)** Venn Diagram showing the overlap of DEG between cohorts. 36 genes of a total of 185 DEG were differentially expressed across all cohorts.

### Supplementary Tables

#### Supplementary Table S1: Top 117 features selected by the DIABLO model

| The top 117 features selected by the DIABLO model | |
|---|---|
| **RNA** | **PROTEIN** |
| RB1 | IL-17C |
| CYBB | IL-2RB |
| VEGFA | Flt3L |
| ZEB1 | ADA |
| CD68 | CRNN |
| TNFRSF9 | VEGFR-3 |
| IL18 | IL-10RA |
| FOXO1 | LIF-R |
| GBP1 | NGF |
| IDO1 | TNFB |
| ID2 | MICA |
| IL3RA | MICB |
| ITGB1 | CD5 |
| PSMB9 | LYPD3 |
| PTPN11 | SPARC |
| CCL5 | CD70 |
| TBP | GZMB |
| SKAP2 | CDCP1 |
| CTSS | EN-RAGE |
| GZMB | SYND1 |
| GRAP2 | HGF |
| NKG7 | WFDC2 |
| PVR | |
| CD226 | |
| DDX58 | |
| BRCA2 | |
| C1QB | |
| IL10 | |
| IL12A | |
| CD53 | |
| CCR2 | |
| TNFRSF17 | |
| CD38 | |
| CD52 | |
| ITGAE | |
| MAPK1 | |
| CDKN3 | |
| IKZF1 | |
| TNFAIP8 | |
| SAMHD1 | |
| TLR7 | |
| HLA-DRA | |

| |
|---|
| SLAMF7 |
| HAVCR2 |
| BTLA |
| IL15 |
| SELL |
| TFRC |
| CD19 |
| FCRLA |
| CXCR2 |
| TNFRSF9 |
| AIF1 |
| CD79A |
| HLA-DQA1 |
| MIF |
| IRF4 |
| NT5E |
| IFITM1 |
| TNFSF10 |
| CTSS |
| CCR4 |
| IRS1 |
| IGSF6 |
| IFITM2 |
| TNFSF13B |
| KIR2DL1 |
| KREMEN1 |
| HLA-DPA1 |
| LMNA |
| ARG1 |
| CD4 |
| CA4 |
| IL2RB |
| ZAP70 |
| IL10RA |
| CCR7 |
| MYC |
| TCF7 |
| CSF1R |
| ITK |
| CD6 |
| SIT100 |
| CD8B |

**Supplementary Databases**

**Supplementary Database S1: DEG between cohorts for Day 1 samples only.**

**Supplementary Database S2: Top GO and Kegg Pathway Functional enrichments in the top features contributing to the classification performance of the final DIABLO model along the first component.**

**Supplementary Database S3: Top GO Pathway Functional enrichments in the top features contributing to the classification performance of the final DIABLO model along the second component.**
